# Complete genome sequence and Benzophenone-3 mineralisation potential of *Rhodococcus* sp. USK10, a bacterium isolated from riverbank sediment

**DOI:** 10.1101/2021.11.04.467327

**Authors:** Joseph Donald Martin, Urse Scheel Krüger, Athanasios Zervas, Morten Dencker Schostag, Tue Kjærgaard Nielsen, Jens Aamand, Lars Hestbjerg Hansen, Lea Ellegaard-Jensen

## Abstract

Benzophenone-3 (BP3) is an organic UV filter whose presence in the aquatic environment has been linked to detrimental developmental impacts in aquatic organisms such as coral and fish. The genus *Rhodococcus* has been extensively studied and is known for possessing large genomes housing genes for biodegradation of a wide range of compounds, including aromatic carbons. Here, we present the genome sequence of *Rhodococcus* sp. USK10, which was isolated from Chinese riverbank sediment and is capable of utilising BP3 as the sole carbon source, resulting in full BP3 mineralisation. The genome consisted of 9,870,030 bp in 3 replicons, a G+C content of 67.2%, and 9,722 coding DNA sequences (CDSs). Annotation of the genome revealed that 179 of these CDSs are involved in metabolism of aromatic carbons. The complete genome of *Rhodococcus* sp. USK10 is the first complete, annotated genome sequence of a Benzophenone-3 degrading bacterium. Through radiolabelling, it is also the first bacterium proven to mineralise Benzophenone-3. Due to the widespread environmental prevalence of Benzophenone-3, coupled to its adverse impact on aquatic organisms, this characterisation provides an integral first step in better understanding the environmentally relevant degradation pathway of the commonly used UV filter. Given USK10’s ability to completely mineralise Benzophenone-3, it could prove to be a suitable candidate for bioremediation application.

## 1. Introduction

Benzophenone-3 (BP3; 2-hydroxy-4-methoxybenzophenone; Oxybenzone) is an organic UV filter typically used in personal care products to protect the skin from harmful solar radiation. Organic UV filters have an aromatic chemical structure that allows for the absorption and stabilisation of both UVA (315-400 nm) and UVB (280-315 nm) radiation [1]. BP3 has been implemented as an active ingredient in sunscreens, cosmetics, and plastic products for decades, and is still one of the most commonly used UV filters worldwide. BP3 has been detected in surface waters, sediments and organisms within various environments including remote areas such as seawater of the Polar Regions [2,3]. Elevated concentrations of BP3 in the aquatic environment have been reported to result in adverse effects in aquatic organisms, such as deterioration of coral reefs and impaired reproduction potential in fish [4–6,1]. These detrimental factors have caused the use of BP3 containing sunscreens to be banned on the coasts of several countries, including the United States (Hawaii, U.S. Virgin Islands), Mexico, and Palau [1,4,7]. The chemical characteristics of BP3, and many other organic UV filters, is a cause of concern due to their high lipophilicity allowing for them to easily bioaccumulate in aquatic organisms and even in the body fluids of humans [2,8]. In addition, BP3 may also act as an endocrine disruptor in humans, influencing birth weight and gestational age [9]. The presence of BP3 in the aquatic environment worldwide begs the question of its persistence and, therefore, it is important to further research the biodegradation potential of BP3 facilitated by microorganisms found in natural environments.

In this study, we isolated and characterized the genome of *Rhodococcus* sp. USK10, to provide additional evidence of the genetic background of this BP3 mineralising bacterium. Currently, only two other bacterial strains, *Methylophilus* sp. strain FP-6 [10] and *Sphingomonas wittichii* strain BP14P [11], have been reported capable of degrading BP3. The phylogenetic characterisation of these strains was however solely based on 16S rRNA gene sequences and their genetic make-up was not investigated. Furthermore, both strains were hypothesized to be able to mineralise BP3, without, however, confirming it experimentally.

Here, we present the first complete and annotated genome of a BP3 degrader found in nature, including a potential linear megaplasmid and a smaller circular plasmid. Strain USK10 shows increased number of genes involved in catalyzing aromatic compounds compared to related *Rhodococcus* strains, which may indicate that it is a specialist strain. In addition, we present experimental data that prove the biodegradation of BP3 by *Rhodococcus* sp. USK10, when incubated in liquid media without any other carbon source.

## 2. Materials and Methods

### 2.1 Isolation of Rhodococcus sp. USK10

Strain USK10 was isolated from enrichment cultures originating from a Chinese riverbank sediment (GPS coordinates 25.569611, 119.781000). The sediment is characterised as unpolluted, having no known prior exposure to BP3. In short, the sediment was implemented into a series of enrichment cultures using radiolabeled BP3 to assess degradation potential followed by a series of streak plating using BP3 enriched agar plates as the sole carbon source. Single colonies were picked and further asses for BP3 mineralisation potential and later characterised, one of which being USK10.

### 2.2 BP3 Biodegradation experiment

Precultures for the mineralisation experiment were grown on R2B media supplemented with 100ppm BP3. After incubation at 20°C in the dark on an orbital shaker (120rpm) for 3 days, extracts were centrifuged (12,000*g* x 5 minutes), washed twice, and resuspended in Difco™ Bushnell-Hass Broth (BHB). The mineralisation experiment was conducted in triplicate with each microcosm containing 5 mL of BHB with BP3 as the sole carbon source. Each USK10 microcosm had approximately 1.4×10^8^ cells, while the abiotic control had no cells. The initial BP3 concentration of each microcosm was 10 mg L^−1^, including [benzene-^14^C(U)]-labeled BP3 (Moravek Biochemicals Inc.; Brea, California, USA) amounting to 7055 DPM. The flasks further contained a 2 mL glass tube with 1 mL 1M NaOH serving as a basetrap to capture the evolved ^14^CO_2_ during BP3 mineralisation. The microcosms were incubated in the dark at 20°C and sampled once a day for 10 days. At each sampling time point, the NaOH was removed, replaced, and transferred to a plastic scintillation vial containing 10 mL of OptPhase HiSafe 3 scintillation cocktail (PerkinElmer, Waltman, MA, USA). All vials were counted for 10min using a Tri-Carb 2810 TR liquid scintillation analyzer (PerkinElmer, Waltman, MA, USA).

### 2.3 DNA Extraction and library preparation

High Molecular Weight DNA was extracted from USK10 grown on R2B liquid media. Prior to DNA extraction, strain purity was confirmed via streak plating on agar plates containing BP3 at a concentration of 250 ppm. DNA extractions were conducted using the Genomic Mini AX Bacteria kit (A&A Biotechnology, Gdynia, Poland). After extraction, the DNA was cleaned and concentrated using the Genomic DNA Clean & Concentrator kit (Zymo Research, Irvine, CA, USA) to remove any impurities that may have been present in the extracts. Concentration and quality of the DNA extracts were measured using Qubit 2.0 Fluorometer with the 1x DS DNA Assay (Invitrogen, Carlsbad, CA, USA) and NanoDrop Spectrophotometer ND-1000 (Thermo Fisher Scientific, Walther, MA, USA), respectively. For Illumina sequencing an Illumina Nextera XT library was prepared for paired-end sequencing on an Illumina NextSeq550 (Illumina Inc., San Diego, CA, USA) according to the manufacturer’s protocol. For Oxford Nanopore sequencing, a library was prepared using the Rapid Sequencing kit (SQK-RBK004) according to the manufacturer’s instructions. Sequencing was performed on a MinION (Oxford Nanopore Technologies, Oxford, UK) with a FLO-MIN106 flow cell, controlled using MinKnow (19.10.1).

### 2.4 Bioinformatics analyses

Sequencing adapters for Illumina reads were trimmed with Trim Galore (0.6.4) (https://github.com/FelixKrueger/TrimGalore). Raw Nanopore fast5 reads were basecalled with GPU-Guppy (3.2.6+afc8e14). A long-read only assembly was created using Raven (1.2.2) [12] and subsequently polished with the Unicycler polish module from the Unicycler assembler (0.4.8) [13], which applies long-read polishing with Racon [14] and short-read polishing with Pilon [15]. The completeness of the genomes was verified by mapping to reference using the Illumina and Nanopore reads with BBmap [16] and Minimap2 [17] under default settings, implemented in Geneious Prime v2020.2.4 (Biomatters). Plasmid sequences were classified using MOB-suite (3.0.0) [18]. The assembled draft genome was annotated using Rapid Annotation using Subsystem Technology (RAST), an online prokaryotic genome annotation platform [19]. Genome completeness was evaluated using BUSCO v5.2.2 using the bacteria_odb10 lineage and “genome” mode [20]. For 16S phylogenetic tree construction, 16S rRNA gene sequences of strains related to USK10 were retrieved by BLAST (https://blast.ncbi.nlm.nih.gov/Blast.cgi). The 16S rRNA gene sequences were collected and aligned with MAAFT [21] under default settings in Geneious Prime v2020.2.4. The alignment was subsequently used for the construction of a 16S rRNA based phylogenetic tree using RAxML [22] in Geneious Prime, specifically the Rapid Bootstrapping and search for best-scoring ML tree algorithm with 100 iterations. Whole genome-based phylogenetic analysis was conducted using the Genome Taxonomy Database (GTDB) [23,24]. The classify workflow (classify_wf) of the Genome Taxonomy Database Toolkit (GTDB-Tk) was used to determined USK10’s taxonomic assignment [25]. The workflow produced a list of genomes similar to that of USK10 along with ANI scores for comparison purposes. Those genome assemblies were retrieved via NCBI and implemented in the lineage workflow (lingeage_wf) of CheckM to assess the similarities of their core genomes [26]. The alignment produced via CheckM was uploaded to Geneious Prime. A phylogenetic tree was created using RAxML, which utilised the GTR GAMMA nucleotide model under the “Rapid Bootstrapping and search for besting-scoring ML tree” algorithm with 100 boostraps replicated.

### 2.5 Data availability

The genome and plasmid sequence of *Rhodococcus* sp. USK10 has been deposited in GenBank under the accession numbers CP076046-CP076048.

## 3. Results and Discussion

### 3.1 BP3 degradation potential of Rhodococcus sp. USK10

BP3 mineralization potential of *Rhodococcus* sp. USK10 was evaluated by measuring released carbon dioxide originating from labelled BP3 added as sole carbon source in a liquid medium microcosm. Figure 1 depicts the complete mineralisation of BP3 by strain USK10. USK10 starts to mineralise BP3 following a two days lag-phase. On the 10th day of the experiment, cumulatively 52.7% of the initial ^14^C label was collected in the form of ^14^CO_2_ and complete mineralisation was assumed. The remaining labelled carbon fraction has likely been incorporated into construction of cellular biomass or metabolites [27]. Comparatively, Lui and colleagues [28] studied biodegradation of BP3 in activated sludge microcosms, focusing on the biodegradation under various redox conditions. They reported that BP3 was completely biodegraded within 42 days of incubation. However, the half-lives of BP3 were observed to be relatively shorter at approximately 4-11 days. *Rhodococcus* sp. USK10 demonstrates the ability to mineralise BP3 within 10 days under aerobic conditions. Furthermore, degradation of BP3 has been shown in water via the UV/H_2_O_2_ and UV/persulfate (UV/PS) reactions, but also using persulfate, metal ions, PbO/TiO_2_ and Sb_2_O_3_/TiO_2_ and other chemicals [29–31]. However, these solutions are not considered “green solutions”.

**Figure 1.**
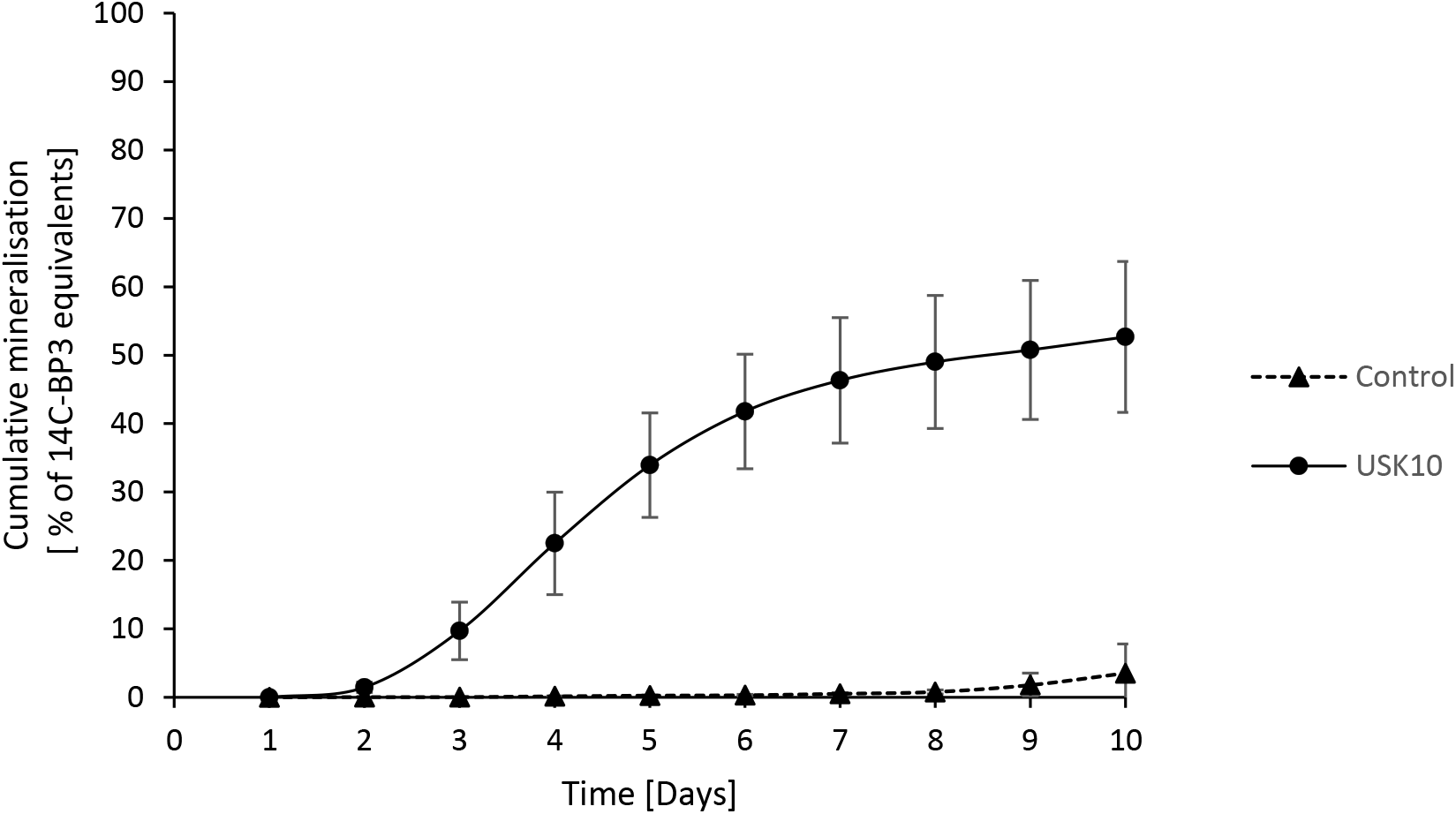
Cumulative mineralisation of BP3 by strain USK10 in pure culture and an abiotic control over ten days. Mean values and standard deviation based on three replicates are shown for ^14^CO_2_ production relative to the initial amount of ^14^C-BP3 added (^14^C_0_).

### 3.2 Genome analysis

The complete genome sequence of *Rhodococcus* sp. USK10 is composed of three replicons with a total assembly of 9,870,030 bp and a G+C content of 67.2%. The chromosome is 8,396,788bp (G+C content 67.6%), while the two mobilisable plasmids are 1,355,759 bp (linear, G+C content 64.6%) and 117,483 (circular, GC content 66%). The genomic map of the chromosome is presented in Figure 2. The circularity of the 3 replicons was verified by mapping-to-reference runs using the Illumina and Nanopore reads in Geneious Prime. For the chromosome and the small plasmid, these were successful. For the larger plasmid, manual forcing of circularity in Geneious Prime and subsequent mapping-to-reference yielded negative results for both the Illumina and Nanopore reads. *Rhodococcus* spp. as well as other Actinobacteria (e.g. *Micrococcus* spp. [32]) are known for having large linear plasmids housing genes coding for degradation potential [33,34]. The assembly is of high quality as revealed by BUSCO analysis (123 complete BUSCOs / 120 single copy and 3 double BUSCOs / 1 fragmented BUSCO / 99,2% coverage of bacteria_odb10).

**Figure 2.**
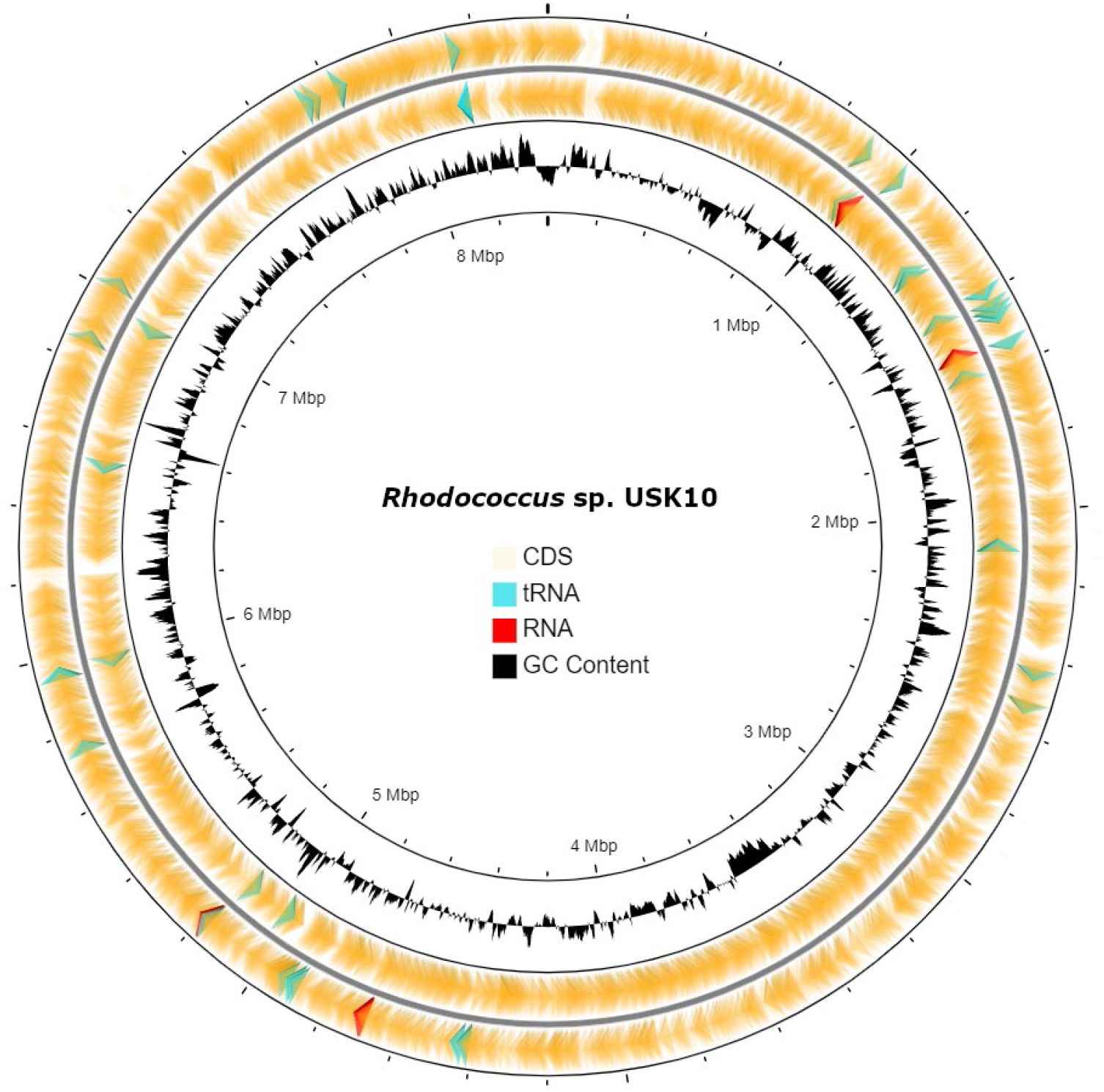
Circular map of *Rhodococcus* sp. USK10 chromosome. The two outer rings represent the coding sequences of the chromosome; the outermost being the forward strand and the innermost being the reverse strand. The inner most ring represents GC content. The G+C content of the chromosome is 67.2%. Created using CGview Server [*35*].

### 3.3 Phylogenetic placement of Rhodococcus sp. USK10

The phylogenetic analysis of both the 16S rRNA gene sequences and whole genome showed that USK10 is well supported within the *Rhodococcus* genus. Based on 16s rRNA gene sequences, *R. wratislaviensis* DSM 44107 and *R. koreensis* DNP505 are the closest relatives of USK10, having pairwise identities of 99.5% and 99.2%, respectively. Figure 3 presents the phylogenetic tree based on 16S rRNA gene sequences.

**Figure 3.**
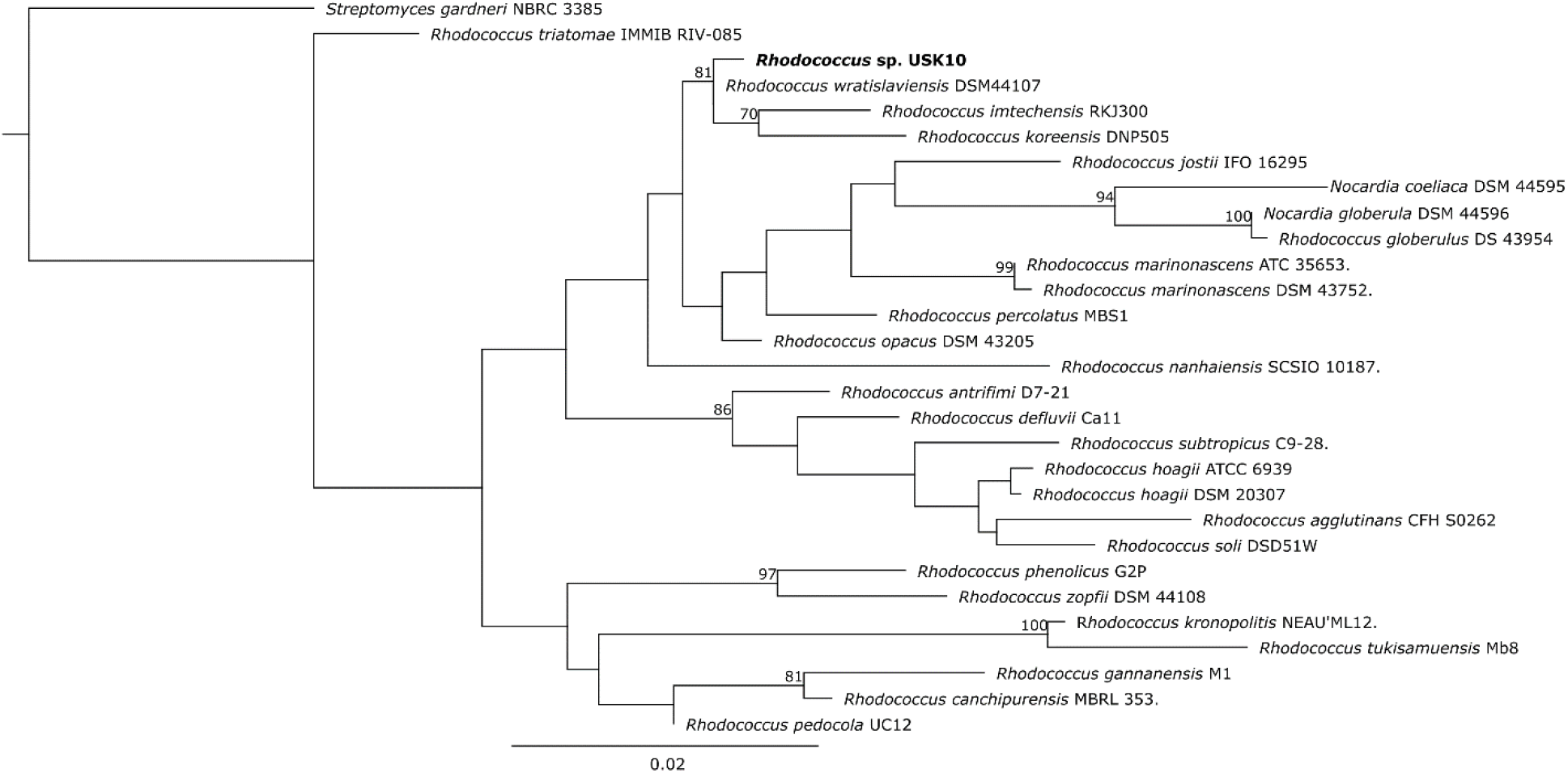
A phylogenetic tree based on 16S rRNA gene sequences showing the position of USK10 in relation to other *Rhodococcus* species and related genera of Actinobacteria. The nucleotide sequences were obtained from NCBI and aligned using the MAAFT alignment plugin via Geneious Prime 2020.2.4. The tree was constructed by RAxML (version 8.2.11) in Geneious Prime 2020.2.4. Node numbers denote bootstrap support values above 60%. The nucleotide module used was GTR GAMMA and the algorithm used was “Rapid Bootstrapping and search for the best-scoring ML tree” with 100 replicates. Bar, 0.02 substitutions per nucleotide position.

For whole genome analysis, GTDB-Tk classified the bacterial genomes based on phylogeny of 120 marker genes and ANI [25]. The ANI scores of each genome in relation to USK10 are presented jointly with the CheckM whole genome tree in Figure 4. From this analysis, the Rhodococcus strain determined to be the most related to USK10 was *Rhodococcus* sp. NCIMB 12038 with an ANI score of 95. This boarders the species demarcation threshold. The next most related bacterium was *Rhodococcus koreensis* DSM 44489 which had an ANI score of 94.92. Considering the limited number of available *Rhodococcus* genomes, the exact ANI threshold for species affiliation is not certain. It has been seen on other bacterial groups (e.g. the *Bacillus cereus* group [36], the genus *Serratia* [37]), that this threshold ranges between 92 and 96%. Based on the topology of both the 16S rRNA sequences and the whole genome comparison, USK10 can be definitely placed and is well supported within the *Rhodococcus* genus. Additional characterisation analyses, such chemotaxonomic and biochemical assays, which are outside the scope of this study, would need to be conducted to assign proper taxonomic assessment to strain USK10 as well as strain NCIMB 12038 to be entirely confident.

**Figure 4.**
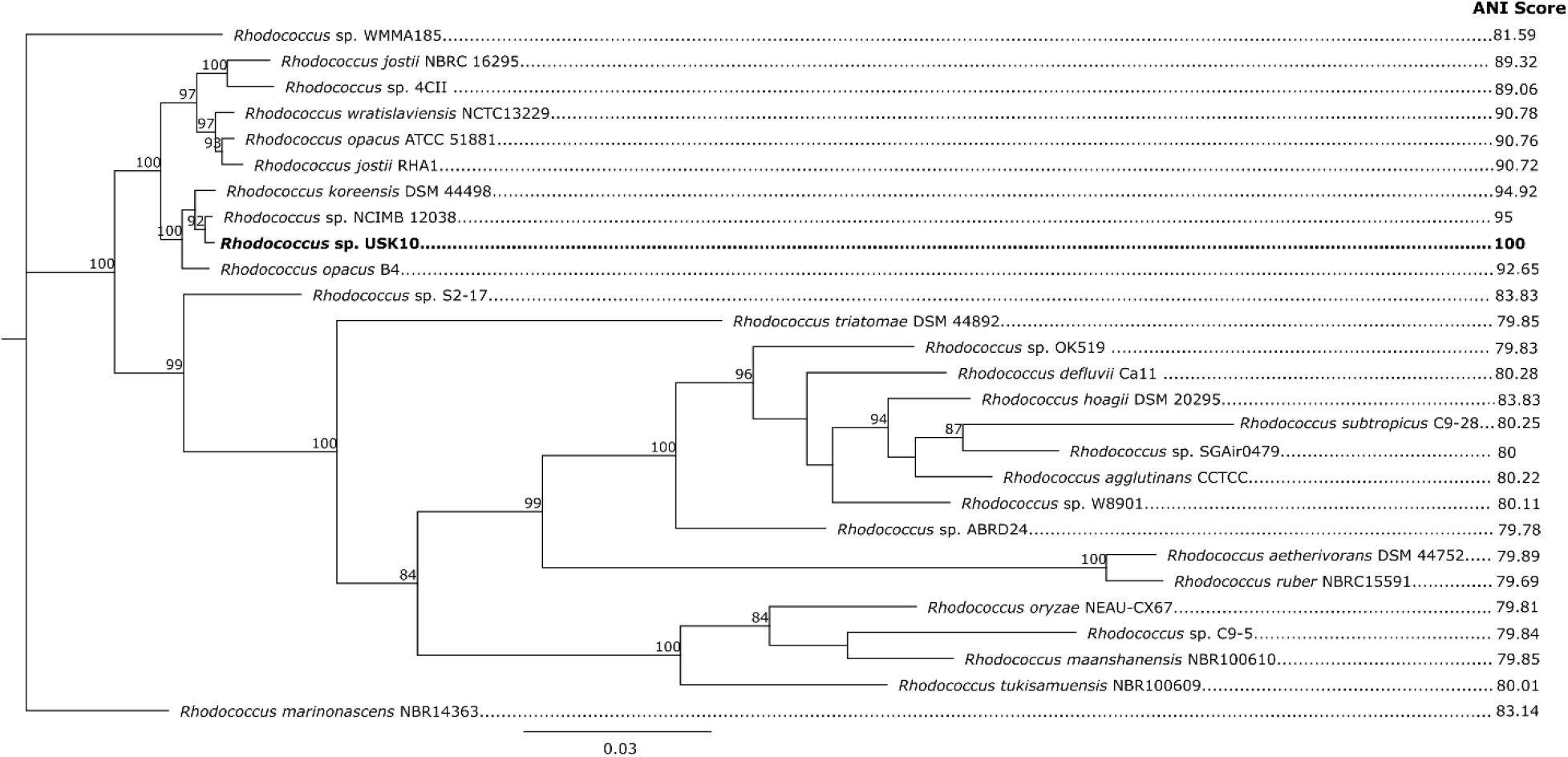
Phylogenetic tree constructed around the position of USK10 based on whole genome sequences using CheckM alignment. The genome sequences were obtained via the NCBI assembly database. The tree was constructed by RAxML (version 8.2.11) in Geneious Prime 2.0. The nucleotide substitution model used was GTR GAMMA and the algorithm used was “Rapid Bootstrapping and search for the best-scoring ML tree” with 100 replicates. Node numbers denote bootstrap support values above 80%. Score values on the right indicate ANI scores obtained via whole genome comparison of USK10 using the GTDB-Tk classify workflow. The distance scale indicates 0.03 substitutions per nucleotide position.

### 3.4 Genome annotation

The annotated genome contains a total of 9,722 CDSs along with 61 RNA encoding genes. RAST was able to provide a general overview of the biological features within the genome, achieving a subsystem coverage of 39% of the annotated genes, 3,817 subsystem feature counts. Of those counts, 179 were responsible for metabolism of aromatic compounds, some of which are likely involved in the degradation process of BP3. Out of the six enzyme classes, five are present in the coding sequences, which involve the metabolism of aromatic compounds (26 hydrolases, 5 isomerases, 15 lyases, and 63 oxidoreductases,). Additionally, 13 transfer proteins involved in the degradation process were identified. The remainder of the CDSs annotated for involvement in metabolism of aromatic compounds, 2 were part of the PcA regulatory protein PcAR family and 39 part of the Transcriptional regulator IclR family. Both these protein families have been well documented to be involved in the degradation of aromatic carbons and present in other *Rhodococcus* species [38]. In *Sphingomonas wittichii* RW1 and DC-6, the first step in the degradation of aromatic compounds is performed by a dioxygenase gene located on a megaplasmid. USK10 bears 6 dioxygenase on its linear megaplasmid, 1 on the small circular plasmid and 65 dioxygenase on its chromosome. Interestingly, the 1 dioxygenase on the small plasmid (3-carboxyethylcatechol 2,3-dioxygenase) is placed next to a FAD-binding monooxygenase (PheA/TfdB family, similar to 2,4-dichlorophenol 6-monooxygenase) which is involved in the degradation of another phenolic compound, 2,4-dichlrophenol. As another alternative, hydroxylases have been suggested to be implemented in the first step of BP3 biodegradation. USK10 possesses 11 hydroxylases on its megaplasmid, 1 on its small plasmid, and 26 on its genome. The megaplasmid contains 65 oxidoreductases that may also play a role in USK10’s biodegradation potential of BP3. Further exploitation of the *Rhodococcus* sp. USK10 genome, and that of other degraders, could lead to more confident identification of potential genes and processes involved in the biodegradation of BP3. Transcriptome sequencing and potentially proteomics analysis of BP3 degrading bacteria may also illuminate the involved genes and pathways in BP3 and other aromatic compounds degradation.

In this regard, *Rhodococcus* sp. USK10 has the potential to be used in large scale efforts to clean BP3-contaminated water sustainably. A key point to be investigated is the survivability and persistence of USK10 in mixed cultures. Other efforts to use microbes for biodegradation of xenobiotics have revealed a plethora of factors that may impact the effect of such approaches that need to be investigated and addressed accordingly [39].

## Acknowledgements

This project was funded by Aarhus University Research Foundation starting grant (AUFF-E-2017-7-21) and Rural Water and Food Security (PI RURAL), the European Commission (Contract No. PI/2017/382/-112).

## Author contributions

Conceptualization: Joseph Donald Martin, Urse Scheel Krüger; Methodology: Joseph Donald Martin, Athanasios Zervas, Urse Scheel Krüger, Tue Kjærgaard Nielsen; Validation: Joseph Donald Martin, Urse Scheel Krüger; Resources: Jens Aamand, Lars Hestbjerg Hansen, Lea Ellegaard-Jensen; Writing: Joseph Donald Martin, Athanasios Zervas, Urse Scheel Krüger, Tue Kjærgaard Nielsen; Review and Editing: Joseph Donald Martin, Athanasios Zervas, Urse Scheel Krüger, Morten Dencker Schostag, Tue Kjærgaard Nielsen, Jens Aamand, Lars Hestbjerg Hansen, Lea Ellegaard-Jensen; Supervision: Lea Ellegaard-Jensen, Morten Dencker Schostag, Jens Aamand, Lars Hestbjerg Hansen.

## References

1. He, T.; Tsui, M.M.P.; Tan, C.J.; Ng, K.Y.; Guo, F.W.; Wang, L.H.; Chen, T.H.; Fan, T.Y.; Lam, P.K.S.; Murphy, M.B. Comparative Toxicities of Four Benzophenone Ultraviolet Filters to Two Life Stages of Two Coral Species. Science of The Total Environment 2019, 651, 2391–2399, doi:10.1016/j.scitotenv.2018.10.148.

2. Tsui, M.M.P.; Leung, H.W.; Wai, T.-C.; Yamashita, N.; Taniyasu, S.; Liu, W.; Lam, P.K.S.; Murphy, M.B. Occurrence, Distribution and Ecological Risk Assessment of Multiple Classes of UV Filters in Surface Waters from Different Countries. Water Research 2014, 67, 55–65, doi:10.1016/j.watres.2014.09.013.

3. Emnet, P.; Gaw, S.; Northcott, G.; Storey, B.; Graham, L. Personal Care Products and Steroid Hormones in the Antarctic Coastal Environment Associated with Two Antarctic Research Stations, McMurdo Station and Scott Base. Environmental Research 2015, 136, 331–342, doi:10.1016/j.envres.2014.10.019.

4. Downs, C.A.; Kramarsky-Winter, E.; Segal, R.; Fauth, J.; Knutson, S.; Bronstein, O.; Ciner, F.R.; Jeger, R.; Lichtenfeld, Y.; Woodley, C.M.; et al. Toxicopathological Effects of the Sunscreen UV Filter, Oxybenzone (Benzophenone-3), on Coral Planulae and Cultured Primary Cells and Its Environmental Contamination in Hawaii and the U.S. Virgin Islands. Arch Environ Contam Toxicol 2016, 70, 265–288, doi:10.1007/s00244-015-0227-7.

5. Balázs, A.; Krifaton, C.; Orosz, I.; Szoboszlay, S.; Kovács, R.; Csenki, Z.; Urbányi, B.; Kriszt, B. Hormonal Activity, Cytotoxicity and Developmental Toxicity of UV Filters. Ecotoxicology and Environmental Safety 2016, 131, 45–53, doi:10.1016/j.ecoenv.2016.04.037.

6. DiNardo, J.C.; Downs, C.A. Dermatological and Environmental Toxicological Impact of the Sunscreen Ingredient Oxybenzone/Benzophenone-3. Journal of Cosmetic Dermatology 2018, 17, 15–19, doi:10.1111/jocd.12449.

7. Fitt, W.K.; Hofmann, D.K. The Effects of the UV-Blocker Oxybenzone (Benzophenone-3) on Planulae Swimming and Metamorphosis of the Scyphozoans Cassiopea Xamachana and Cassiopea Frondosa. Oceans 2020, 1, 174–180, doi:10.3390/oceans1040013.

8. Kim, S.; Choi, K. Occurrences, Toxicities, and Ecological Risks of Benzophenone-3, a Common Component of Organic Sunscreen Products: A Mini-Review. Environment International 2014, 70, 143–157, doi:10.1016/j.envint.2014.05.015.

9. Ghazipura, M.; McGowan, R.; Arslan, A.; Hossain, T. Exposure to Benzophenone-3 and Reproductive Toxicity: A Systematic Review of Human and Animal Studies. Reproductive Toxicology 2017, 73, 175–183, doi:10.1016/j.reprotox.2017.08.015.

10. Jin, C.; Geng, Z.; Pang, X.; Zhang, Y.; Wang, G.; Ji, J.; Li, X.; Guan, C. Isolation and Characterization of a Novel Benzophenone-3-Degrading Bacterium Methylophilus Sp. Strain FP-6. Ecotoxicology and Environmental Safety 2019, 186, 109780, doi:10.1016/j.ecoenv.2019.109780.

11. Fagervold, S.K.; Rohée, C.; Rodrigues, A.M.S.; Stien, D.; Lebaron, P. Efficient Degradation of the Organic UV Filter Benzophenone-3 by Sphingomonas Wittichii Strain BP14P Isolated from WWTP Sludge. Science of The Total Environment 2021, 758, 143674, doi:10.1016/j.scitotenv.2020.143674.

12. Vaser, R.; Šikić, M. Raven: A de Novo Genome Assembler for Long Reads; 2021; p. 2020.08.07.242461;

13. Wick, R.R.; Judd, L.M.; Gorrie, C.L.; Holt, K.E. Unicycler: Resolving Bacterial Genome Assemblies from Short and Long Sequencing Reads. PLOS Computational Biology 2017, 13, e1005595, doi:10.1371/journal.pcbi.1005595.

14. Vaser, R.; Sović, I.; Nagarajan, N.; Šikić, M. Fast and Accurate de Novo Genome Assembly from Long Uncorrected Reads. Genome Res. 2017, 27, 737–746, doi:10.1101/gr.214270.116.

15. Walker, B.J.; Abeel, T.; Shea, T.; Priest, M.; Abouelliel, A.; Sakthikumar, S.; Cuomo, C.A.; Zeng, Q.; Wortman, J.; Young, S.K.; et al. Pilon: An Integrated Tool for Comprehensive Microbial Variant Detection and Genome Assembly Improvement. PLOS ONE 2014, 9, e112963, doi:10.1371/journal.pone.0112963.

16. Bushnell, B.; Rood, J.; Singer, E. BBMerge – Accurate Paired Shotgun Read Merging via Overlap. PLOS ONE 2017, 12, e0185056, doi:10.1371/journal.pone.0185056.

17. Li, H. Minimap2: Pairwise Alignment for Nucleotide Sequences. Bioinformatics 2018, 34, 3094–3100, doi:10.1093/bioinformatics/bty191.

18. Robertson, J.; Nash, J.H.E. MOB-Suite: Software Tools for Clustering, Reconstruction and Typing of Plasmids from Draft Assemblies. Microb Genom 2018, 4, e000206, doi:10.1099/mgen.0.000206.

19. Aziz, R.K.; Bartels, D.; Best, A.A.; DeJongh, M.; Disz, T.; Edwards, R.A.; Formsma, K.; Gerdes, S.; Glass, E.M.; Kubal, M.; et al. The RAST Server: Rapid Annotations Using Subsystems Technology. BMC Genomics 2008, 9, 75, doi:10.1186/1471-2164-9-75.

20. Simão, F.A.; Waterhouse, R.M.; Ioannidis, P.; Kriventseva, E.V.; Zdobnov, E.M. BUSCO: Assessing Genome Assembly and Annotation Completeness with Single-Copy Orthologs. Bioinformatics 2015, 31, 3210–3212, doi:10.1093/bioinformatics/btv351.

21. Katoh, K.; Standley, D.M. MAFFT Multiple Sequence Alignment Software Version 7: Improvements in Performance and Usability. Molecular Biology and Evolution 2013, 30, 772–780, doi:10.1093/molbev/mst010.

22. Stamatakis, A. RAxML-VI-HPC: Maximum Likelihood-Based Phylogenetic Analyses with Thousands of Taxa and Mixed Models. Bioinformatics 2006, 22, 2688–2690, doi:10.1093/bioinformatics/btl446.

23. Parks, D.H.; Chuvochina, M.; Waite, D.W.; Rinke, C.; Skarshewski, A.; Chaumeil, P.-A.; Hugenholtz, P. A Standardized Bacterial Taxonomy Based on Genome Phylogeny Substantially Revises the Tree of Life. Nat Biotechnol 2018, 36, 996–1004, doi:10.1038/nbt.4229.

24. Parks, D.H.; Chuvochina, M.; Chaumeil, P.-A.; Rinke, C.; Mussig, A.J.; Hugenholtz, P. A Complete Domain-to-Species Taxonomy for Bacteria and Archaea. Nat Biotechnol 2020, 38, 1079–1086, doi:10.1038/s41587-020-0501-8.

25. Chaumeil, P.-A.; Mussig, A.J.; Hugenholtz, P.; Parks, D.H. GTDB-Tk: A Toolkit to Classify Genomes with the Genome Taxonomy Database. Bioinformatics 2020, 36, 1925–1927, doi:10.1093/bioinformatics/btz848.

26. Parks, D.H.; Imelfort, M.; Skennerton, C.T.; Hugenholtz, P.; Tyson, G.W. CheckM: Assessing the Quality of Microbial Genomes Recovered from Isolates, Single Cells, and Metagenomes. Genome Res. 2015, 25, 1043–1055, doi:10.1101/gr.186072.114.

27. Bouchez, M.; Blanchet, D.; Vandecasteele, J.-P. The Microbiological Fate of Polycyclic Aromatic Hydrocarbons: Carbon and Oxygen Balances for Bacterial Degradation of Model Compounds. Appl Microbiol Biotechnol 1996, 45, 556–561, doi:10.1007/BF00578471.

28. Liu, Y.-S.; Ying, G.-G.; Shareef, A.; Kookana, R.S. Biodegradation of the Ultraviolet Filter Benzophenone-3 under Different Redox Conditions. Environmental Toxicology and Chemistry 2012, 31, 289–295, doi:10.1002/etc.749.

29. Lee, Y.-M.; Lee, G.; Zoh, K.-D. Benzophenone-3 Degradation via UV/H2O2 and UV/Persulfate Reactions. Journal of Hazardous Materials 2021, 403, 123591, doi:10.1016/j.jhazmat.2020.123591.

30. Pan, X.; Yan, L.; Qu, R.; Wang, Z. Degradation of the UV-Filter Benzophenone-3 in Aqueous Solution Using Persulfate Activated by Heat, Metal Ions and Light. Chemosphere 2018, 196, 95–104, doi:10.1016/j.chemosphere.2017.12.152.

31. Wang, Z.; Deb, A.; Srivastava, V.; Iftekhar, S.; Ambat, I.; Sillanpää, M. Investigation of Textural Properties and Photocatalytic Activity of PbO/TiO2 and Sb2O3/TiO2 towards the Photocatalytic Degradation Benzophenone-3 UV Filter. Separation and Purification Technology 2019, 228, 115763, doi:10.1016/j.seppur.2019.115763.

32. Dib, J.R.; Wagenknecht, M.; Hill, R.T.; Farías, M.E.; Meinhardt, F. First Report of Linear Megaplasmids in the Genus Micrococcus. Plasmid 2010, 63, 40–45, doi:10.1016/j.plasmid.2009.10.001.

33. König, C.; Eulberg, D.; Gröning, J.; Lakner, S.; Seibert, V.; Kaschabek, S.R.; Schlömann, M. A Linear Megaplasmid, P1CP, Carrying the Genes for Chlorocatechol Catabolism of Rhodococcus Opacus 1CP. Microbiology 2004, 150, 3075–3087, doi:10.1099/mic.0.27217-0.

34. Gűrtler, V.; Seviour, R.J. Systematics of Members of the Genus Rhodococcus (Zopf 1891) Emend Goodfellow et al. 1998. In Biology of Rhodococcus; Alvarez, H.M., Ed.; Microbiology Monographs; Springer: Berlin, Heidelberg, 2010; pp. 1–28 ISBN 978-3-642-12937-7.

35. Stothard, P.; Wishart, D.S. Circular Genome Visualization and Exploration Using CGView. Bioinformatics 2005, 21, 537–539, doi:10.1093/bioinformatics/bti054.

36. Zervas, A.; Aggerbeck, M.R.; Allaga, H.; Güzel, M.; Hendriks, M.; Jonuškienė, Ii.; Kedves, O.; Kupeli, A.; Lamovšek, J.; Mülner, P.; et al. Identification and Characterization of 33 Bacillus Cereus Sensu Lato Isolates from Agricultural Fields from Eleven Widely Distributed Countries by Whole Genome Sequencing. Microorganisms 2020, 8, 2028, doi:10.3390/microorganisms8122028.

37. Serratia Inhibens Sp. Nov., a New Antifungal Species Isolated from Potato (Solanum Tuberosum) | Microbiology Society Available online: https://www.microbiologyresearch.org/content/journal/ijsem/10.1099/ijsem.0.004270?crawler=true (accessed on 1 November 2021).

38. Tropel, D.; van der Meer, J.R. Bacterial Transcriptional Regulators for Degradation Pathways of Aromatic Compounds. Microbiology and Molecular Biology Reviews 2004, 68, 474–500, doi:10.1128/MMBR.68.3.474-500.2004.

39. Hylling, O.; Nikbakht Fini, M.; Ellegaard-Jensen, L.; Muff, J.; Madsen, H.T.; Aamand, J.; Hansen, L.H. A Novel Hybrid Concept for Implementation in Drinking Water Treatment Targets Micropollutant Removal by Combining Membrane Filtration with Biodegradation. Science of The Total Environment 2019, 694, 133710, doi:10.1016/j.scitotenv.2019.133710.

